# Nitrogen resorption efficiency in autumn leaves correlated with chlorophyll resorption, not with anthocyanin production

**DOI:** 10.1101/2021.05.12.443938

**Authors:** Ines Pena-Novas, Marco Archetti

## Abstract

A prominent hypothesis for the adaptive value of anthocyanin production in the autumn leaves of some species of trees is that anthocyanins protect leaves from photooxidative stress at low temperatures, allowing a better resorption of nutrients – in particular, nitrogen – before leaf fall. While there is evidence that anthocyanins enable photoprotection, it is not clear whether this translates to improved nitrogen resorption and how this can explain inter-specific variation in autumn colours. A recent comparative analysis showed no correlation between temperature and anthocyanin production across species but did not analyse nitrogen content and nitrogen resorption efficiency. Here we provide this comparison by comparing the nitrogen content of mature and senescent leaves and their autumn colours in 55 species of trees. We find no correlation between the presence of anthocyanins and the efficiency of nitrogen resorption. We find, instead, that nitrogen resorption is more efficient in species with yellow autumn colours, pointing to chlorophyll resorption, rather than anthocyanin synthesis, as the main determinant of nitrogen resorption efficiency. Hence our results do not corroborate the photoprotection hypothesis as an explanation for the evolution of autumn colours.

## Introduction

There is still debate [Renner and Zhoner 2020, Pena-Novas & Archetti 2020a] on the adaptive value of autumn colours. While yellow is understood to be a by-product of the breakdown of chlorophyll that allows the colour of carotenoids to stand out [Lee 2002, Tanaka et al. 2008], the production of anthocyanins that make the leaves of some species of trees and shrubs red in autumn is controversial. Among current hypotheses [Archetti 2009a], a long-standing [Pringsheim 1879] and prominent one is the photoprotection hypothesis [Field al. 2001, Hoch et al. 2003, Lee et al. 2003]: anthocyanins protect leaves from photooxidative stress. Protection is especially important in autumn, when cold temperatures reduce carbon fixation capacity, light is no longer fully used for photosynthesis due to chlorophyll breakdown, and light is more intense due to a thinning canopy [Ougham et al. 2008]. While protecting senescing leaves themselves is not critical for the plant, photoprotection may allow a better resorption of nutrients. Nitrogen, in particular, is often a limiting factor that strongly influences tree growth [Dickson 1989, Cooke et al. 2005, Schlesinger 2009], and trees have evolved multiple adaptations to reabsorb nitrogen seasonally [Van Cleve and Apel 1993]. The photoprotection hypothesis suggests that the adaptive value of anthocyanin synthesis in autumn is to enable a more efficient resorption of nutrients, especially nitrogen, at low temperatures [Field al. 2001, Hoch et al. 2003, Lee et al. 2003].

Anthocyanins can indeed function as antioxidants [Lee 2002, Lee & Gould 2002, Nagata et al. 2003, Kytridis & Manetas 2006] especially at low temperatures [Gould et al. 2018], even though anthocyanins in leaves may be neither ideal nor ideally located to protect against light [Manetas 2006; Duan et al. 2014]. Empirical evidence for the photoprotection hypothesis has come mainly from observations on the photoprotective effects of anthocyanins. While some evidence clearly supports the hypothesis [Manetas et al. 2002, Hughes et al. 2005, 2007, Neill et al. 2002a, Schaberg et al. 2003, Hoch et al. 2003], some does not or is unclear [Burger & Edwards 1996, Feild et al. 2001, Lee et al. 2003, Manetas et al. 2003, Hormaetxe 2005, Karageorgou & Manetas 2006, Kyparissis et al. 2007, Esteban et al. 2008, Neill et al. 2002b, Hughes et al. 2005, Feild et al. 2001, Lee et al. 2003, Nikiforou & Manetas 2010, Nikiforou et al. 2011]. Some previous studies report a difference between red and non-red leaves in nitrogen resorption efficiency [Schaberg et al. 2003, Hoch et al. 2003] but some do not [Field et al. 2001, Duan et al 2014] or are unclear [Lee et al. 2003].

While the photoprotection hypothesis remains plausible [Archetti et al. 2009, Renner and Zhoner 2020, Pena-Novas & Archetti 2020a], evidence that anthocyanins provide photoprotection and enhance nitrogen resorption is necessary but not sufficient to prove that photoprotection is the actual adaptive values of autumn colours. In an ideal experiment, populations would evolve with or without the need for photoprotection and we would observe whether autumn colours arise only in the populations under selection. In other words, the question is not just why autumn colours exist, but why they exist only in some species, and additional evidence must come from comparative phylogenetic analysis.

A recent study [Renner and Zhoner 2020] suggests that there is a higher prevalence of tree species with autumn colours in Eastern North America (ENA) than in Europe because ENA has lower temperatures, and that this supports the photoprotection hypothesis. A re-analysis of that dataset, however, showed that there is no significant difference in the prevalence of red autumn colours between ENA and Europe [Pena-Novas & Archetti 2020a]. More importantly, it would be more appropriate to compare the growing temperatures and autumn colours of individual species, rather than comparing arbitrary floras. A recent analysis [Pena-Novas & Archetti 2020b] comparing the climatic parameters of 237 tree species revealed no significant difference in growing temperature between species with red autumn leaves and species with green or yellow autumn leaves – a result that does not corroborate the photoprotection hypothesis.

A limitation of this analysis, however, is that it did not test whether species with red autumn leaves reabsorb nitrogen more efficiently than species with non-red leaves. This is what we do here, using a comparative analysis of data on nitrogen resorption we gathered from 55 tree species from the same single location.

## Materials and methods

### Autumn colours and nitrogen content

Data were collected at the H.O. Smith Arboretum of Pennsylvania State University (40.805051°N, 77.864088°W). Autumn leaf colour (a categorical variable with three possible states: green, yellow and red) was observed directly by the authors in November 2019 and 2020. Nitrogen content was measured from leaves collected on various days in Autumn 2020 (“Autumn”); on 21 September 2020 (“Summer”); and on 21 September 2019 (“Previous Summer”). We define resorption efficiency as 1-*r*, where *r* is the ratio between the Autumn and Summer (2020) nitrogen content (the lower the ratio, the higher the resorption efficiency). Leaves were collected on the same day for all species in Summer; in Autumn, leaves were collected on different days for different species, when ready for abscission (when it was possible to detach them from the tree by a slight touch). Nitrogen content was measured (by the Agricultural Analytical Services Laboratory of Pennsylvania State University) using CN Elementar analyzers (a Vario Max Cube in 2019; and a Rapid Max N Exceed in 2020). Both instruments use the Dumas combustion method to measure nitrogen. Nitrogen content is reported as percent of leaf material.

### Comparative analysis

Statistical analysis of the differences in nitrogen parameters is reported as the result of a Mann–Whitney U test. To control for phylogeny, we then analyse character correlation using the pairwise comparison method [Read & Nee 1995, Maddison 2000] implemented in the *Mesquite* software [Maddison & Maddison 2018] on a recent phylogeny of trees [Zanne et al. 2014]. The method chooses phylogenetically independent pairs of species (the path between members of a pair, along the branches of the tree, does not touch the path of any other pair) to avoid pseudo-replication, and tests whether a difference in one character (nitrogen parameter) consistently predicts a difference in the second character (autumn colour).

## Results

### Our analysis of nitrogen content in 55 tree species (Figure 1, Table 1) reveals the following results

Nitrogen content values in mature leaves over two consecutive summers are, not surprisingly, correlated; values for the summer and the following autumn are also correlated; nitrogen resorption efficiency is correlated with nitrogen content in autumn but not in summer (Figure 2, Table 2)

**Figure 1.**
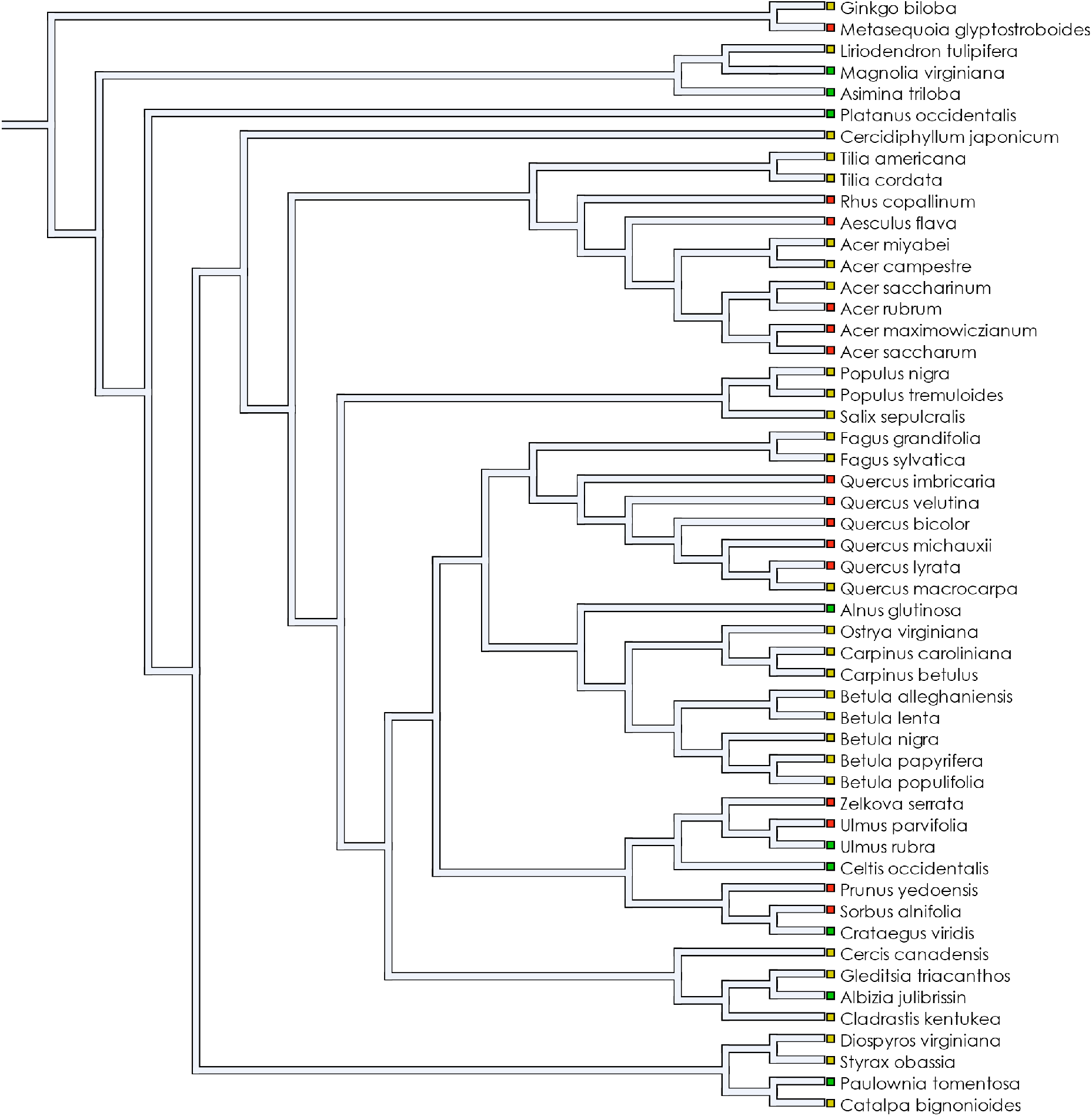
Species used in the analysis. The phylogenetic tree of the species used in the analysis and their autumn colours.

**Figure 2.**
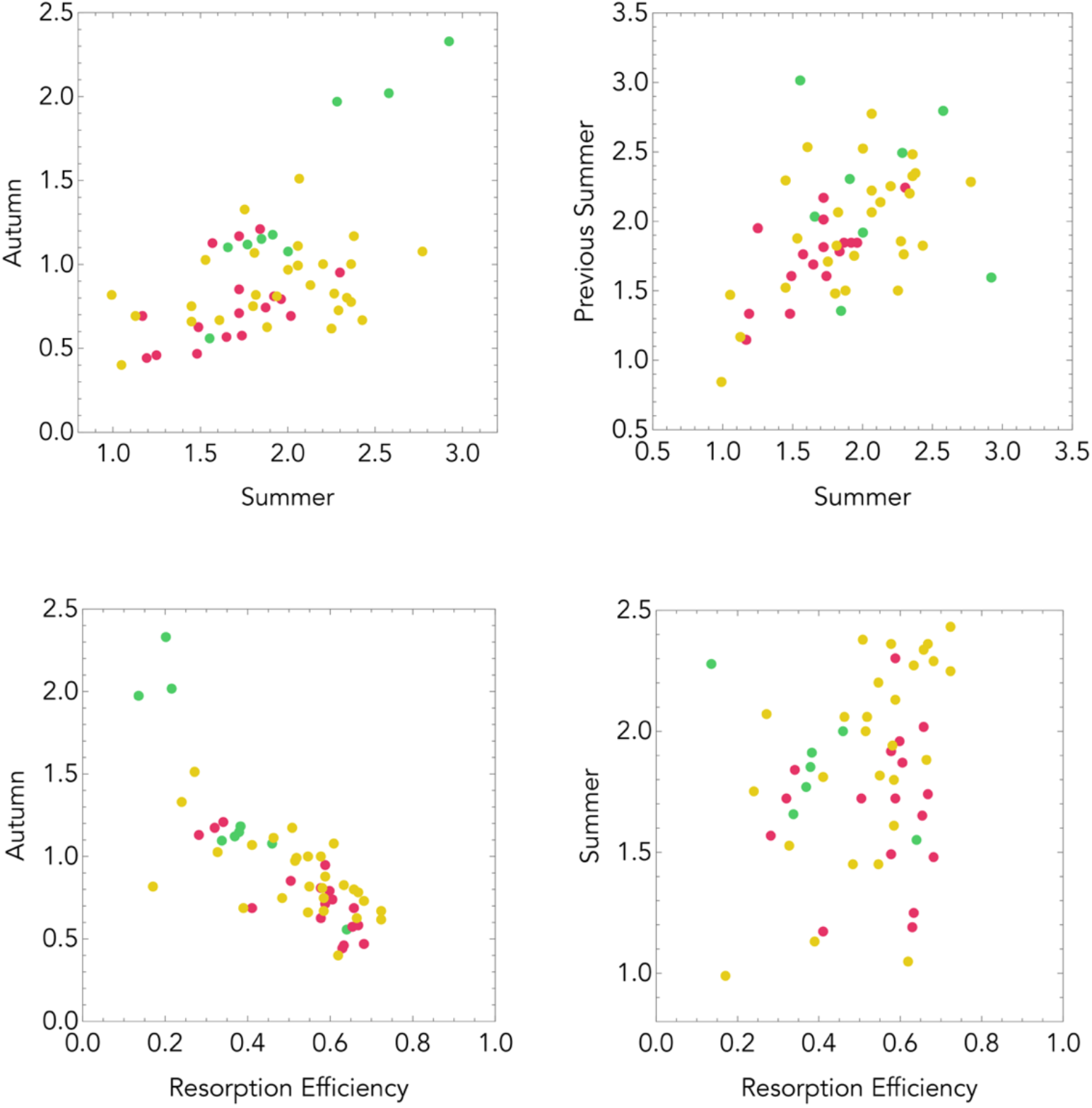
Correlations between nitrogen parameters. Scatter plots of nitrogen content in Autumn, Summer, Previous Summer and Resorption Efficiency.

**Table 1.** Species used in the analysis. For each species, autumn colours, nitrogen content in Autumn 2020, Summer 2020, their ratio, and nitrogen content in the previous Summer (2019) are listed.

| Species | Colour | Autumn | Summer | Ratio | Previous Summer |
| --- | --- | --- | --- | --- | --- |
| *Acer campestre* | Yellow | 1.11 | 2.06 | 0.54 | 2.35 |
| *Acer freemanii* | Red | 1.21 | 1.84 | 0.66 | 1.78 |
| *Acer maximowiczianum* | Red | 0.63 | 1.49 | 0.42 | 1.61 |
| *Acer miyabei* | Yellow | 1.07 | 1.81 | 0.59 | 1.82 |
| *Acer rubrum* | Red | 0.47 | 1.48 | 0.32 | 1.76 |
| *Acer saccharinum* | Yellow | 0.97 | 2.00 | 0.49 | 2.52 |
| *Acer saccharum* | Red | 0.44 | 1.19 | 0.37 | 1.95 |
| *Aesculus flava* | Red | 0.69 | 1.17 | 0.59 | 1.33 |
| *Albizia julibrissin* | Green | 1.97 | 2.28 | 0.87 | 2.49 |
| *Alnus glutinosa* | Green | 2.02 | 2.58 | 0.78 | 3.01 |
| *Asimina triloba* | Green | 1.08 | 2.00 | 0.54 | 1.92 |
| *Betula alleghaniensis* | Yellow | 0.78 | 2.36 | 0.33 | 2.25 |
| *Betula lenta* | Yellow | 1.00 | 2.20 | 0.46 | 2.30 |
| *Betula nigra* | Yellow | 0.88 | 2.13 | 0.41 | 2.14 |
| *Betula papyrifera* | Yellow | 0.73 | 2.29 | 0.32 | 1.76 |
| *Betula populifolia* | Yellow | 0.62 | 2.25 | 0.27 | 2.07 |
| *Carpinus betulus* | Yellow | 0.99 | 2.06 | 0.48 | 2.07 |
| *Carpinus caroliniana* | Yellow | 0.75 | 1.45 | 0.52 | - |
| *Catalpa bignonioides* | Yellow | 1.33 | 1.75 | 0.76 | 1.71 |
| *Celtis occidentalis* | Green | 1.10 | 1.66 | 0.66 | 2.31 |
| *Cercidiphyllum japonicum* | Yellow | 0.40 | 1.05 | 0.38 | 1.47 |
| *Cercis canadensis* | Yellow | 1.17 | 2.38 | 0.49 | 2.48 |
| *Cladrastis kentukea* | Yellow | 0.80 | 2.34 | 0.34 | 2.20 |
| *Crataegus viridis* | Green | 1.15 | 1.85 | 0.62 | 1.36 |
| *Diospyros virginiana* | Yellow | 0.75 | 1.80 | 0.42 | 1.48 |
| *Fagus grandifolia* | Yellow | 0.83 | 2.27 | 0.36 | 2.28 |
| *Fagus sylvatica* | Yellow | 1.03 | 1.53 | 0.67 | 1.88 |
| *Ginkgo biloba* | Yellow | 0.69 | 1.13 | 0.61 | 1.17 |
| *Gleditsia triacanthos* | Yellow | 0.81 | 1.94 | 0.42 | 1.75 |
| *Liriodendron tulipifera* | Yellow | 0.63 | 1.88 | 0.34 | 1.50 |
| *Magnolia acuminata* | Yellow | 0.66 | 1.45 | 0.45 | 1.52 |
| *Magnolia virginiana* | Green | 0.56 | 1.55 | 0.36 | 2.03 |
| *Metasequoia glyptostroboides* | Red | 1.17 | 1.72 | 0.68 | 1.33 |
| *Ostrya virginiana* | Yellow | 0.82 | 1.82 | 0.45 | 1.86 |
| *Paulownia tomentosa* | Green | 2.33 | 2.92 | 0.80 | 2.80 |
| *Platanus occidentalis* | Green | 1.18 | 1.91 | 0.62 | - |
| *Populus nigra* | Yellow | 0.67 | 2.43 | 0.28 | 1.83 |
| *Populus tremuloides* | Yellow | 1.51 | 2.07 | 0.73 | - |
| *Prunus yedoensis* | Red | 0.57 | 1.65 | 0.35 | 2.17 |
| *Quercus bicolor* | Red | 0.74 | 1.87 | 0.40 | 1.85 |
| *Quercus imbricaria* | Red | 0.81 | 1.92 | 0.42 | - |
| *Quercus lyrata* | Red | 0.71 | 1.72 | 0.41 | 1.69 |
| *Quercus macrocarpa* | Yellow | 0.99 | 2.06 | 0.48 | 2.22 |
| *Quercus michauxii* | Red | 0.85 | 1.72 | 0.49 | 1.81 |
| *Quercus muehlenbergii* | Red | 0.79 | 1.96 | 0.40 | 1.85 |
| *Quercus velutina* | Red | 0.69 | 2.02 | 0.34 | 2.01 |
| *Rhus copallinum* | Red | 0.46 | 1.25 | 0.37 | 1.15 |
| *Salix sepulcralis* | Yellow | 0.67 | 1.61 | 0.42 | 2.53 |
| *Sorbus alnifolia* | Red | 0.58 | 1.74 | 0.33 | 1.61 |
| *Styrax obassia* | Yellow | 0.82 | 0.99 | 0.83 | 0.84 |
| *Tilia americana* | Yellow | 1.08 | 2.77 | 0.39 | 2.77 |
| *Tilia cordata* | Yellow | 1.00 | 2.36 | 0.43 | 2.33 |
| *Ulmus parvifolia* | Red | 1.13 | 1.57 | 0.72 | 1.85 |
| *Ulmus rubra* | Green | 1.12 | 1.77 | 0.63 | 1.60 |
| *Zelkova serrata* | Red | 0.95 | 2.30 | 0.41 | 2.24 |

**Table 2.** Correlations between nitrogen parameters. Statistics and significance levels (*P*) for the Spearman Rank test and the Pearson Correlation test (reported only if all the *P*-values resulting from a test for normality are above 0.025) for the correlations between nitrogen content in Summer, Autumn, Previous Summer and Resorption Efficiency. Degrees of freedom are in parenthesis.

|  | Summer,<br>Previous<br>(49) | Summer,<br>Autumn<br>(51) | Summer,<br>Resorption<br>(51) | Autumn,<br>Resorption<br>(51) |
| --- | --- | --- | --- | --- |
| Spearman Rank | 0.67, $P=2\times 10^{-8}$ | 0.46, $P=4\times 10^{-4}$ | 0.05, $P=0.69$ | -0.78, $P=6\times 10^{-14}$ |
| Pearson Correlation | 0.75, $P=2\times 10^{-11}$ | | 0.11, $P=0.41$ | |

When data are not controlled for phylogeny (Figure 3, Table 3), nitrogen content in summer is higher in species with green autumn leaves than in species with red autumn leaves (it is higher in species with yellow autumn leaves than in species with red autumn leaves only in one of the two years examined); nitrogen content in autumn is higher in species with green autumn leaves than in species with red or yellow autumn leaves. Therefore, it appears that nitrogen resorption efficiency is higher in species with red autumn leaves (as predicted by the photoprotection hypothesis). Nitrogen resorption efficiency is, however, also higher in species with yellow autumn leaves than in species with green leaves, which is not predicted by the photoprotection hypothesis; and there is no difference in resorption efficiency between species with red or yellow autumn leaves, which is against the predictions of the photoprotection hypothesis.

**Figure 3.**
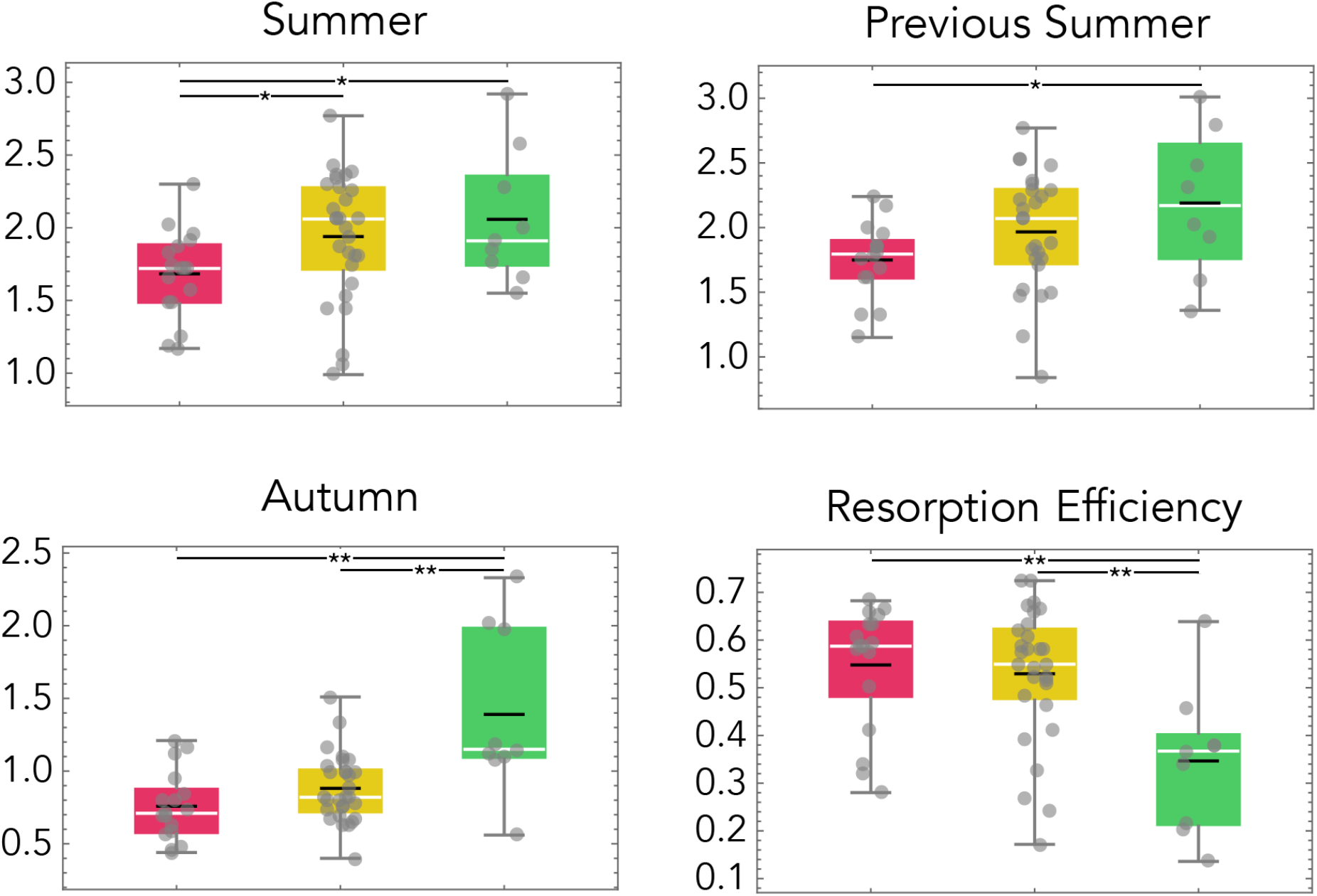
Differences in nitrogen parameters between autumn colours. Species are grouped by autumn colours (red, yellow, green). Box plots show the median (white bar); mean (black bar); 25% and 75% quartiles (coloured areas); upper and lower fences (grey bars); individual data points (grey dots). Asterisks shows significant *P* values in a Mann–Whitney *U* test (\**P*<0.05; \*\**P*<0.01; see Table 3 for the exact values).

**Table 3.** Differences in nitrogen parameters between autumn colours. Statistics and significance levels (*P*) of the Mann–Whitney *U* test for the differences in nitrogen parameters between autumn colours. Asterisks shows significant values (\**P*<0.05; \*\**P*<0.01). See Figure 3 for value distribution.

|  | Red, Yellow |  | Red, Green |  | Green, Yellow |  |
| --- | --- | --- | --- | --- | --- | --- |
| Autumn | 171 | $P=0.090$ | 26 | $P=0.006^{**}$ | 44 | $P=0.003^{**}$ |
| Summer | 144 | $P=0.020^{*}$ | 40 | $P=0.046^{*}$ | 124 | $P=0.823$ |
| Previous Summer | 142 | $P=0.067$ | 32 | $P=0.046^{*}$ | 83 | $P=0.316$ |
| Resorption efficiency | 279 | $P=0.452$ | 128 | $P=0.009^{**}$ | 214 | $P=0.004^{**}$ |

When the same analysis is repeated, more appropriately, controlling for phylogeny (Table 4), the difference in resorption efficiency between red and green species is no longer significant. The difference in resorption efficiency between green and yellow species remains.

**Table 4.** Phylogenetic independent contrast for the differences in nitrogen parameters between autumn colours. Differences in nitrogen content in Autumn, Summer and Resorption efficiency are tested between species grouped by autumn colour. The total number of pairings used in each test is shown in parenthesis. Table entries are the values (or ranges) of the *P* values for the best tail of each pairing. Significant values are in bold.

|  | Red<br>Non red<br>(2) | Red<br>Green<br>(2) | Red<br>Yellow<br>(43) | Green<br>Yellow<br>(74) | Green<br>Non green<br>(2) |
| --- | --- | --- | --- | --- | --- |
| Resorption efficiency | 0.23 | 0.23 | [0.25, 0.69] | **[0.016,0.50]** | [0.06,0.11] |
| Autumn | 0.23 | 0.06 | [0.25, 0.69] | **[0.016,0.34]** | [0.11,0.23] |
| Summer | 0.23 | 0.06 | [0.12, 0.69] | **[0.016,0.50]** | [0.11,0.23] |

Therefore, chlorophyll resorption (non-green autumn leaves; particularly for yellow leaves), rather than anthocyanins production (red leaves) seems to be the major determinant of nitrogen resorption efficiency.

## Discussion

In summary, we found no evidence that red leaves (the presence of anthocyanins) enable a more efficient resorption of nitrogen compared to green or yellow leaves, which is inconsistent with the photoprotection hypothesis. Species with yellow autumn leaves reabsorb nitrogen more efficiently than species with green leaves, a prediction that is not directly relevant for the photoprotection hypothesis, but that points to chlorophyl resorption (which enables the yellow colour of carotenoids to become visible) or a correlated process, rather than anthocyanin production, as the major determinant of nitrogen resorption efficiency.

Recent results on a larger dataset [Pena-Novas & Archetti 2020b] showed that tree species growing at colder temperature are more likely to have yellow, not red, autumn leaves, suggesting that resorption of chlorophyll, but not the production of anthocyanins, might be an adaptation to cold temperatures. While the dataset we use here is smaller, comparing nitrogen resorption efficiency is a direct test of the photoprotection hypothesis, which posits that the adaptive value of red autumn leaves is an efficient resorption of nutrients, particularly nitrogen, enabled by the photoprotective effect of anthocyanins. Photoprotection is known to be more important at lower temperatures – hence the rationale for analysing climate parameters [Pena-Novas & Archetti 2020b] to test indirectly the assumptions of the photoprotection hypothesis. Analysing nitrogen resorption efficiency is a more direct test of the hypothesis – a test that, based on the results we report here, does not support the hypothesis.

We must point out that the power of our analysis is relatively low, given that we use only 55 species. While phylogenetic pairwise comparison avoids pseudoreplication of lineage-specific factors, allowing a proper statistical analysis, it uses only a subset of the data [Felsenstein 1985], hence it has a low power to detect correlations [Grafen & Ridley 1996]. We note, however, that a significant difference was found for yellow and green.

While analysing nitrogen content and nitrogen resorption efficiency is an advance over the use of indirect data such as climate parameters, this is not yet a conclusive test of the hypothesis. The photoprotection hypothesis posits that selection for reabsorbing nitrogen is the driving force behind the evolution of autumn colours. A more refined analysis should control for nitrogen *need*, which is arguably different among species. Hence, species for which nitrogen is not a limiting factor would be under weaker selection for photoprotection and would have weaker selective pressure to evolve autumn colours. One possible way to control for nitrogen requirements would be to compare species with and without nitrogen-fixing bacteria (although a cursory analysis reveals that many tree species *without* nitrogen-fixing bacteria [Wright et al. 2004] do not turn red in autumn [Archetti 2009b]).

While our comparative analysis does not support the photoprotection hypothesis for the evolution of autumn colours, more sophisticated analyses might reveal a correlation that our study was unable to detect. Based on our current results, however, one possible conclusion is that nitrogen resorption depends mainly on the resorption of chlorophyll, or a related process, rather than on the photoprotective effects of anthocyanins.

The results that yellow autumn colours correlated with a more efficient resorption of nitrogen has implications for another hypothesis for the evolution of red autumn colours. Among current hypotheses [Archetti 2009a], the coevolution hypothesis suggests that anthocyanins provide a warning signal to pest insects that migrate to trees in autumn [Archetti 2000; Hamilton & Brown 2001], a type of aposematism [Lev-Yadun & Gould 2007]. The hypothesis has been corroborated by comparative analysis [Hamilton & Brown 2001; Archetti 2009c] and empirical studies showing that insects avoid red leaves [Archetti & Leather 2005, Karageorgou & Manetas 2006, Doring et al. 2009, Archetti 2009c], that the amount of red is positively correlated with chemical defences [Karageorgou et al. 2008, Cooney et al. 2012; Menzies et al. 2015], and that insects grow better on plants with green autumn leaves than on plants with red autumn leaves [Archetti 2009c; Maskato et al. 2014]. While the coevolution hypothesis is agnostic about the evolution of *yellow* autumn leaves, it is known that yellow attracts insects [Holopainen & Peltonen 2002, Wilkinson et al. 2002, Doring et al. 2009], acting as a “supergreen” stimulus. If yellow evolved as an adaptation to low temperatures or to enable an efficient resorption of nitrogen, it could have served a pre-adaptation for the subsequent evolution of red, as trees with yellow leaves would be under stronger selective pressure to repel insects (which are attracted by yellow) by evolving chemical defense and correlated warning signals (anthocyanins in red leaves).

Why species with yellow autumn leaves reabsorb nitrogen more efficiently remains to be invertigated. Merely reabsorbing chlorophyll cannot be responsible for a significant difference in nitrogen resorption efficiency, since chlorophyll typically accounts for only a small fraction of total leaf nitrogen [Hörtensteiner 2006], while most of it is in the form of proteins in the leaves. Seasonal resorption of nitrogen involves degradation of these proteins to transportable forms, its transport from senescing leaves and conversion to storage proteins [Cooke and Weih 2005, Sample and Babst 2019]. While protein degradation begins early in the season [Hörtensteiner and Feller 2002, Cooke and Weih 2005], nitrogen is exported from the leaf with a 1–2-month delay [Sample and Babst 2019], at the same time as chlorophyll degradation [Lee et al. 2003]. The result that yellow autumn colours are correlated with efficient nitrogen resorption, suggests that some process linked to chlorophyll resorption is the main determinant of nitrogen resorption efficiency.

The result that red autumn colours are not, suggests that the contribution of anthocyanin to nitrogen resorption is negligible, a result that does not corroborate the photoprotection hypothesis.

## Notes

### Competing Interest Statement

The authors have declared no competing interest.

## References

Archetti M (2000) The origin of autumn colours by coevolution. J. Theor. Biol. 205, 625–630

Archetti M (2009a) Phylogenetic analysis reveals a scattered distribution of autumn colours. Annals of Botany 103:703–713.

Archetti M (2009b) Classification of hypotheses for the evolution of autumn colours. Oikos 118:328–333.

Archetti M (2009c) Evidence from the domestication of apple for the maintenance of autumn colours by coevolution. Proc. Royal Soc. B 276: 2575–2580

Archetti M, et al. (2009) Unravelling the evolution of autumn colours: an interdisciplinary approach. Trends Ecol. Evol. 24: 166–173

Burger, J., Edwards, G.E. (1996). Photosynthetic efficiency, and photodamage by UV and visible radiation in red versus green leaf Coleus varieties. Plant and Cell Physiology 37, 395–399.

Cooke, J.E.K., Martin, T.A., Davis, J.M. (2005) Short-term physiological and developmental responses to nitrogen availability in hybrid poplar, New Phytol. 167: 41–52

Cooke, J.E.K., Weih, M. (2005) Nitrogen storage and seasonal nitrogen cycling in *Populus:* bridging molecular physiology and ecophysiology. New Phytologist 167: 19–30.

Dickson, R.E. (1989) Carbon and nitrogen allocation in trees. Ann Sci For 46:S631–S647.

Duan B, Paquette A, Juneau P, Brisson J, Fontaine B, Berninger F (2014) Nitrogen resorption in Acer platanoides and Acer saccharum: influence of light exposure and leaf pigmentation. Acta Physiol Plant 36:3039–3050.

Esteban R, Fernández-Marín B, Becerril JM, García-Plazaola JI (2008) Photoprotective implications of leaf variegation in *E. dens-canis* L. and *P. officinalis* L. J. Plant Physiol. 165:1255–1263

Felsenstein J (1985). Phylogenies and comparative method. Am. Nat. 125, 1–15.

Field TS, Lee DW, Holbrook NM (2001) Why leaves turn red in autumn. The role of anthocyanins in senescing leaves of red-osier dogwood. Plant Physiol 127: 566–574

Gould KS, Jay-Allemand C, Logan BA, Baissac Y, Bidel LPR (2018) When are foliar anthocyanins useful to plants? Re-evaluation of the photoprotection hypothesis using *Arabidopsis thaliana* mutants that differ in anthocyanin accumulation. Environmental and Experimental Botany 154: 11–12.

Grafen A, Ridley M (1996). Statistical tests for discrete cross-species data. J. Theor. Biol. 183, 255–267.

Hamilton WD, Brown SP (2001) Autumn tree colours as a handicap signal. Proc. Royal Soc. B 268: 1489–1493.

Hoch WA et al. (2003) Resorption protection. Anthocyanins facilitate nutrient recovery in autumn by shielding leaves from potentially damaging light levels. Plant Physiol 133:1296–1305

Holopainen JK, Peltonen P (2002) Bright autumn colours of deciduous trees attract aphids: nutrient retranslocation hypothesis. Oikos 99: 184–188.

Hormaetxe K, Becerril JM, Fleck I, Pintó M, García-Plazaola JI (2005) Functional role of red (retro)-carotenoids as passive light filters in the leaves of *Buxus sempervirens* L.: increased protection of photosynthetic tissues? J. Exp. Bot 56:2629–2636

Hörtensteiner S. (2006). Chlorophyll degradation during senescence. Annual Review of Plant Biology 57: 55–77.

Hughes NM, Morley CB, Smith WK (2007) The coordination of anthocyanin decline and photosynthetic maturation in developing leaves of three deciduous tree species. New Phytol. 175:75–685

Hughes NM, Neufeld HS, Burkey KO (2005) Functional role of anthocyanins in high-light winter leaves of the evergreen herb. Galax urceolata. New Phytol. 168:575–587

Karageorgou P, Buschmann C, Manetas Y (2008) Red leaf colour as a warning signal against insect herbivory: honest or mimetic? Flora 203:648–652.

Karageorgou P, Manetas Y (2006) The importance of being red when young, anthocyanins and the protection of young leaves of *Quercus coccifera* from insect herbivory and excess light. Tree Physiol. 26:613–621

Kyparissis A, Grammatikopoulos G, Manetas Y. (2007) Leaf morphological and physiological adjustments to the spectrally selective shade imposed by anthocyanins in *Prunus cerasifera*. Tree Physiol. 6:849–857

Kytridis VP, Manetas Y (2006) Mesophyll versus epidermal anthocyanins as potential in vivo antioxidants: evidence linking the putative antioxidant role to the proximity of the oxy-radical source. J. Exp. Bot. 57:2203–2210

Lee DW (2002) Anthocyanins in autumn leaf senescence. Adv. Bot. Res. 37, 147–165

Lee DW, Gould KS (2002) Anthocyanins in leaves and other vegetative organs: an introduction. Adv. Bot. Res. 37:1–16

Lee DW, O’Keefe J, Holbrook NM, Field TS (2003) Pigment dynamics and autumn leaf senescence in a New England deciduous forest, eastern USA. Ecol. Res. 18:677–694

Maddison WP (2000) Testing Character Correlation using Pairwise Comparisons on a Phylogeny. J. Theor. Biol. (2000) 202, 195–204

Maddison WP, Maddison DR (2018) Mesquite: a modular system for evolutionary analysis. Version 3.51 http://www.mesquiteproject.org

Manetas Y (2006) Why some leaves are anthocyanic and why most anthocyanic leaves are red? Flora 201: 163–177.

Manetas Y, Drinia A, Petropoulou Y (2002) High contents of anthocyanins in young leaves are correlated to low pools of xanthophyll cycle components and low risk of photoinhibition. Photosynthetica 40:349–354

Manetas Y, Petropoulou Y, Psaras GK, Drinia A (2003) Exposed red (anthocyanic) leaves of *Quercus coccifera* display shade characteristics. Funct. Plant Biol. 30:265–270

Nagata T, Todoriki S, Masumizu T, Suda I, Furuta S, Du Z, Kikuchi S (2003) Levels of active oxygen species are controlled by ascorbic acid and anthocyanin in *Arabidopsis*. J. Agric. Food Chem. 51:2992–2999

Neill SO, Gould KS, Kilmartin PA, Mitchell KA, Markham KR (2002a) Antioxidant activities of red versus green leaves in *Elatostema rugosum*. Plant Cell Environ. 25:537–549

Neill SO, Gould KS, Kilmartin PA, Mitchell KA, Markham KR (2002b) Antioxidant capacities of green and cyanic leaves in the sun species *Quintinia serrata*. Funct. Plant Biol. 29:1437–1443.

Nikiforou, C., Nikolopoulos, D., Manetas, Y. (2011) The winter-red-leaf syndrome in *Pistacia lentiscus:* evidence that the anthocyanic phenotype suffers from nitrogen deficiency, low carboxylation efficiency and high risk of photoinhibition. J Plant Physiol. 168: 2184–7.

Nikiforou, C., Manetas, Y. (2010) Strength of winter-leaf redness as an indicator of stress vulnerable individuals in *Pistacia lentiscus*. Flora 205: 424–427.

Ougham, H., Hörtensteiner, S., Armstead, I., Donnison, I., King, I., Thomas, H., & Mur, L. (2008). The control of chlorophyll catabolism and the status of yellowing as a biomarker of leaf senescence. Plant Biology, 10: 4–14.

Pena-Novas, I., Archetti, M. (2020a) Biogeography and evidence for adaptive explanations of autumn colours. New Phytologist. 228: 809–813.

Pena-Novas, I., Archetti, M. (2020b) A comparative analysis of the photoprotection hypothesis for the evolution of autumn colours. J Evol Biol 33: 1669–1676.

Pringsheim N (1879) *Ueber Lichtwirkung und Chlorophyllfunction in der Planze*. Jahrbuch fuer Wissenschaftliche Botanik. Boratrager, Berlin.

Read AF, Nee S (1995) Inference from binary comparative data. J. Theor. Biol. 173, 99–108

Sample, R., Babst, B.A. (2019) Timing of nitrogen resorption-related processes during fall senescence in Southern Oak species, Forest Science 65: 245–249,

Schaberg PG, Van Den Berg AK, Murakami PF, Shane JB, Donnelly JR (2003) Factors influencing red expression in the autumn foliage of sugar maple trees. Tree Physiol. 23:325–333

Schlesinger, W.H. (2009) On the fate of anthropogenic nitrogen. Proc. Natl. Acad. Sci. USA 106: 203–208.

Tanaka Y et al. (2008) Biosynthesis of plant pigments: anthocyanins, betalains and carotenoids. Plant J. 54, 733–749

Van Cleve, B., Apel, K. (1993) Induction by nitrogen and low temperature of storage-protein synthesis in poplar trees exposed to long days. Planta 189: 157–160.

Wright IJ, Reich PB, Westoby M, Ackerly DD, Baruch Z, Bongers F, Cavender-Bares J, Chapin T, Cornelissen JH, Diemer M, Flexas J, Garnier E, Groom PK, Gulias J, Hikosaka K, Lamont BB, Lee T, Lee W, Lusk C, Midgley JJ, Navas ML, Niinemets U, Oleksyn J, Osada N, Poorter H, Poot P, Prior L, Pyankov VI, Roumet C, Thomas SC, Tjoelker MG, Veneklaas EJ, Villar R. (2004) The worldwide leaf economics spectrum. Nature 428: 821–827.

Zanne AE, Tank DC, Cornwell WK, Eastman JM, Smith SA, FitzJohn RG, McGlinn DJ, O’Meara BC, Moles AT, Reich PB, Royer DL, Soltis DE, Stevens PF, Westoby M, Wright IJ, Aarssen L, Bertin RI, Calaminus A, Govaerts R, Hemmings F, Leishman MR, Oleksyn J, Soltis PS, Swenson NG, Warman L, Beaulieu JM (2014) Three keys to the radiation of angiosperms into freezing environments. Nature 506(7486): 89–92.

